# Residue communities reveal evolutionary signatures of *γδ* T-Cell receptor

**DOI:** 10.1101/2022.12.29.522230

**Authors:** Ngaam J. Cheung, Si-Yu Huang

## Abstract

Naturally co-occurring amino acids, term coevolution, in a protein family play a significant role in both protein engineering and folding, and it is expanding in recent years from the studies of the effects of single-site mutations to the complete re-design of a protein and its folding, especially in three-dimensional structure prediction. Here, to better characterize such coevolving interactions, we *in silico* decipher evolutionary couplings from massive homologous sequences using spectral analysis to capture signatures that are important for specific molecular interactions and binding activities. We implement the present approach on the G7 gamma delta T-cell receptor to identify functionally important residues that contribute to its highly distinct binding mode. The analysis indicates the evolutionary signatures (highly ordered networks of coupled amino acids, termed residue communities) of the protein confirm previously identified functional sites that are relevant to dock the receptor underneath the major histocompatibility complex class I-related protein-1 (MR1) antigen presenting groove. Moreover, we analyze the correlation of inter-residue contacts with the activation states of receptors and show that contact patterns closely correlating with activation indeed coincide with these sites. The theoretical results demonstrate our method provides an alternative path towards bridging protein sequence with its function at residue-level without requiring its tertiary structure or highly accurate measurement of its biological activities *in vivo/vitro*.

## 1 Introduction

Naturally coevolution between residues in a protein and among residues from different proteins are the basis of protein function in most biological processes under selective pressure. The pressure also shapes various distribution of amino acids in different protein families, and particularly, conservation of an amino acid at a position in a protein family is functionally relevant under constraints presented by nature on its evolution. As a result, the distribution of amino acids in proteins is not random^1^, and it encodes information during evolution for specifying protein fold^2,3^ and function^4^. Conservation has long been used to infer residues of a protein that are most likely to be functionally significant^5,6^. Moreover, coevolution information hidden in protein sequences has arisen as an important constraint for accurately predicting protein 3D structures using physics-informed methods^7^ and deep learning-based methods^8,9^.

As advanced by the availability of large scale biological sequences and high-performance computational methods, we are able to mine higher order patterns (knowledge) from those sequences, either site-independent conservation or coevolution between pairwise residues at different sites on a protein sequence. Many advances have been developed to evaluate the importance of residues in proteins to derive precise information for protein 3D structures^7,10^ and function^4^. For example, statistical coupling analysis (SCA) analyzes multiple homologous sequences to find the correlation information among residues with sequence conservation, and those coevolving residues are termed “sectors” that are relevant to biochemical properties of the PDZ domain protein^3^. The evolutionary couplings (ECs) model (also known as direct coupling analysis, DCA), an approach that measures the strength of the direct relationship between pairwise residues at two sites of a biological sequence by distinguishing true coevolution couplings from the noisy correlations^7,11^, has been presented to mine the evolutionary sequence record not only for protein residue contacts^12,13^ and protein-protein interaction networks^14,15^, but also for RNA structure prediction^16,17^.

The gamma delta T-cell receptors (*γδ*TCRs) play an important biological role in human anti-infection immunity^18^, anti-tumor immunity^19^, tissue homeostasis^20^, and mucosal immunity^21^. Different from alpha beta T-cell receptors (*αβ*TCRs), the *γδ*TCRs do not require antigen processing and major-histocompatibility-complex (MHC) presentation of peptide epitopes^22^. The *γδ*TCRs may recognize CD1d–*α*-galactosylceramide (*α*-GalCer)^23^, and they also bind soluble or membrane proteins^22^. In the past decades, research effort has been devoted to advance our understanding of molecular mechanisms and functional role the *γδ*TCRs^22–28^, and many profiling has reveled that the *γδ*TCRs may have specific residues that can be activated by antigenic molecules^27^. Studies have shown that major histocompatibility complex (MHC) molecules or MHC-like molecules can be recognized in an antigen independent manner^24–26^. Since high-throughput screens still remains labor intensive, computational analyses have been advanced to identify residues that are functionally interacted between the *γδ*TCRs and their partners. Although it is significant to decipher the evolutionary information that holds biological relevance from sequences of the *γδ*TCRs for a molecular basis, identifying those coevolving residues from these sequences alone still remains challenging in molecular biology. One long-standing approach for predicting the interactions between them is to search for coevolving amino acids at pairwise sites in aligned multiple homologous sequences. In particular, how the evolutionary information specifies protein’s biological properties and its function in organisms and a comprehensive signatures of proteins have not been completely investigated for the *γδ*TCRs. Challenges of deciphering fingerprints on the *γδ*TCRs that are relevant to their functional sites, e.g., difficult to predict allosteric sites, binding pockets, docking partners, also seemingly grows with neither experimental measurements nor structures. This fact has greatly hindered progress towards an efficient means of accurately recognizing physical interactions between either proteins or proteins and small molecules in the *γδ*TCRs, in which these interactions are critical in either biological processes or drug developments.

Here we present a physics-informed approach to investigate the evolutionary signatures from natural evolution, enabling us to quantitatively measure the energetic landscape of protein from sequences alone. The method takes advantage of the massive biological sequences to quantify the evolutionary effects of perturbations. In the present work, we apply the approach to analyze the residue communities on the G7 *γδ*TCR^29^, and we identify two communities of residues are crucial for recognizing binding molecules and provide an evolutionary bias for how the *γδ*TCR interacts with antigen presented by antigen-presenting molecules. Comparisons to known experimental measurements demonstrate the the accuracy of identifying signatures in the G7 *γδ*TCR.

## 2 Results

### 2.1 Evolutionary information from protein sequences

Studies have revealed that some TCRs can bind underneath the MR1 antigen-binding cleft instead of recognizing the presented antigen^29–31^, but it remains unclear how the binding strategy evolves in the TCRs. Among the *γδ*TCRs, the human G7 *γδ*TCR undergoes different binding strategies that many *αβ*TCR adopt. Recently, Le Nours et al. identified that human *γδ*TCs exhibit auto-reactivity to the MR1^29^ using strikingly different binding modes involving complete rearrangement docking topology (Fig. 1a, right). The sharply distinct mode is unexpected in the known *αβ*TCR, e.g., the MAIT *αβ*TCR binds on the MR1 platform (Fig. 1a, left), suggesting that the G7 *γδ*TCR has a fundamentally different binding mode that is not adopted by many *αβ*TCR^29^. The MAIT executes two essential binding actions using one site, but the G7 *γδ*TCRs docks itself to the MR1 using an unlikely manner. In the present study, we mined the biological sequence data for clues of the molecular mechanisms and performed theoretical analyses on models of the MAIT *αβ*TCR-MR1-5-OP-RU and the G7 *γδ*TCR-MR1-5-OP-RU ternary complexes. Accordingly, we firstly analyzed the conservation of each amino acid of the MR1 from both the MAIT *αβ*TCR-MR1-5-OP-RU and the G7 *γδ*TCR-MR1-5-OP-RU ternary complex^29^ (Fig. 1a) using the relative entropy^32^ (see Eq. (2) in Method). Site-independent conservation computed from multiple aligned sequences (MSAs) of a protein family is a general means to quantitatively access the importance of each amino acid. Generally, highly conserved residues are most likely to be at functional sites of the protein. The relative entropy at each site was computed from the two MR1 MSAs (Fig. 1b). The distributions of conservation at the position of each protein suggest they share similar patterns of conservation in the MR1s, and the degree of conservation of each site was mapped to their tertiary structures (Fig. 1b). Whereas using the conservation information, we cannot find distinguishable differences between the two MR1 proteins, that is, in this case, the different binding modes cannot be inferred from either structural analysis or conservation of residues.

**Figure 1.**
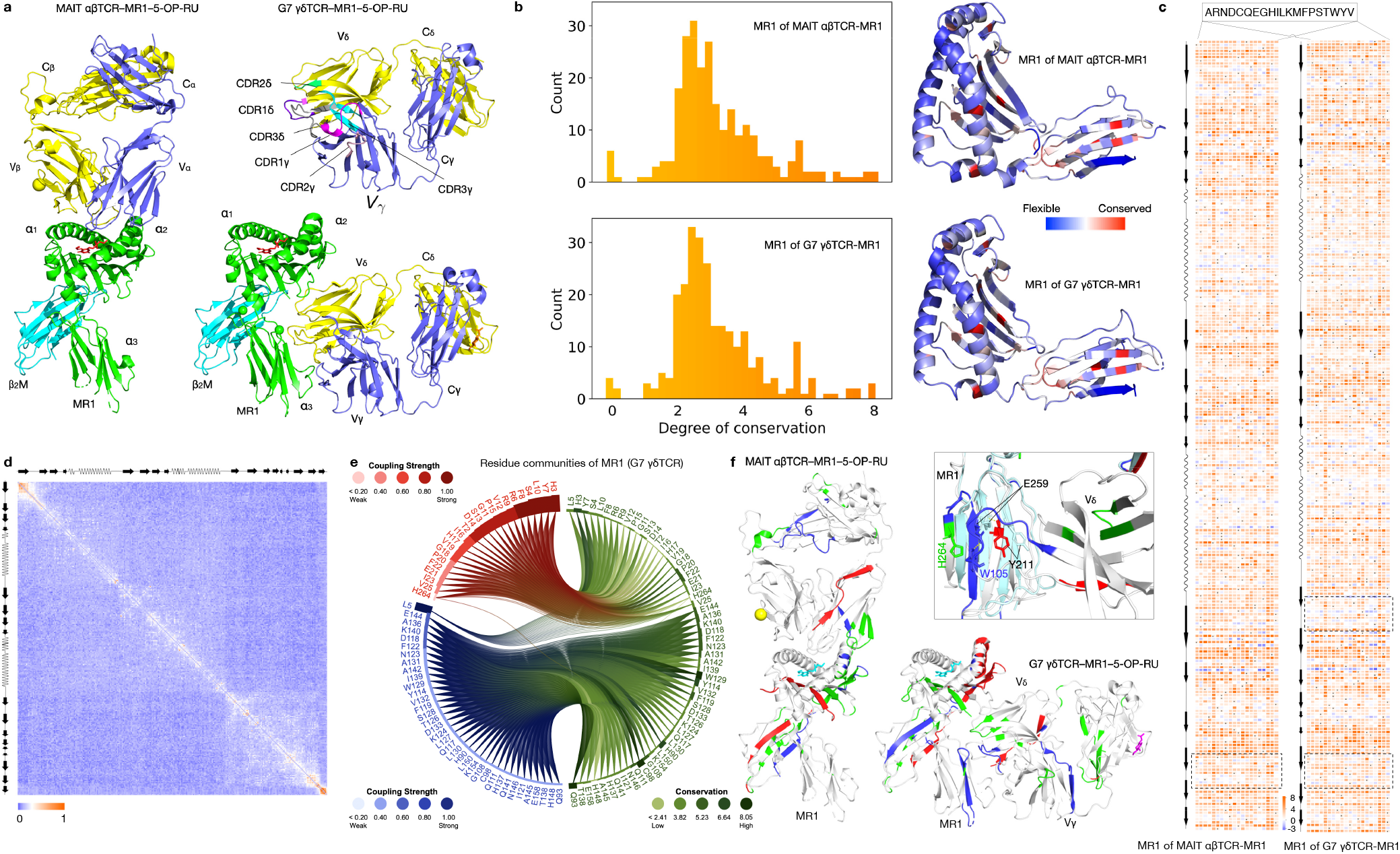
Residue communities mapped on the the *γδ*TCRs. (a) Overview of the MAIT *αβ*TCR-MR1-5-OP-RU (PDB code: 4NQC) and the G7 *γδ*TCR-MR1-5-OP-RU (PDB code: 6MWR) complexes. The MAIT *αβ*TCR-MR1-5-OP-RU complex (left): the MR1 in green, *β*_2_-microglobulin (*β*_2_M) in cyan, 5-OP-RU in red, *α*-chain in blue, and *β* -chain in yellow. The G7 *γ* det TCR-MR1-5-OP-RU complex (right): the MR1 in green, *β*_2_M in cyan, 5-OP-RU in red, *δ* -chain in blue, and *γ*-chain in yellow. On the top-right, CDR1*γ* in magenta, CDR2*γ* in, CDR3*γ* in cyan, CDR1*δ* in purple, CDR2*δ* in green, and CDR3*δ* in gray. (b) Conservation information of the MR1 proteins in the two complexes. Histograms show the distributions of degree of conservation at each sites (left). The scores of conservation are mapped on the structures of the MR1 (right). (c) Perturbation of amino acids in the MR1s of the the MAIT *αβ*TCR-MR1 (left) and the G7 *γδ*TCR-MR1 (right) complexes. The energy-like score of each substitution at a single site is computed using the sequence statistical potential. Dotted squares include slightly different patterns. (d) Evolutionary couplings of the MR1 in the G7 *γδ*TCR-MR1. (e) Residue communities. The top two residue communities with couplings strength are in red and blue, respectively, and their conservation is shown in green. (f) Residue communities mapped on structures. The residues in different communities are colored in red, blue, green, and the remain residues are all in gray. The comparison between *α*_3_ regions of the MR1 proteins is shown on top right.

Next, we followed the present evolutionary spectrum approach to quantitatively examine the distinct binding modes. Leveraging the sequence statistical potential (see Method), we performed comparison of energy-like scores between the two MR1 proteins, the residues close to the C-terminus (*α*_3_) form more but shorter *β* -strands in the G7 *γδ*TCR-MR1 than those in the MAIT *αβ*TCR-MR1, in which the flexibility of the residues is stronger than those of much inflexible. Evolutionary couplings between pairwise residues were computed from the same aligned multiple sequences of the MR1 protein in the *γδ*TCR-MR1 complex, and there are two clear patterns that represent *α*_1_-*α*_2_ and *α*_3_ (Fig. 1d). The coupled signals show that residues of *α*_3_ have weaker co-evolving connections with those of *α*_1_-*α*_2_, nevertheless, they exhibit stronger signals inside of *α*_3_ and have clearly physical connections. The evolutionary couplings reflects that *α*_3_ may play a critical role for contacting *γδ*TCRs, that agrees with finding in the the G7 *γδ*TCR-MR1 complex primarily using the *α*_3_ domain of MR1 for its reactivity^29^.

To better understand the evolution of the distinct binding modes in the G7 *γδ*TCR, we calculated the coupling relationship among residues from sequences alone using evolutionary coupling analysis^33^ (Fig. 1d). The approach utilizes a multiple sequence alignment across different species to estimate couplings between pairwise amino acids. We generated multiple sequence alignments by searching each protein in the G7 *γδ*TCR against the UniRef90 database, from which we estimated the couplings using the maximum likelihood method. The analysis indicates that the MAIT and HLA TCRs hold similar evolutionary signatures, while the G7 *γδ*TCR has a different pattern from those in MAIT and HLA TCRs (Fig. 1). Further, we performed spectral analysis on the evolutionary couplings for detecting understandable signatures, termed residue communities that hold functional information of residues. The communities have clearly interpretable functional roles (Fig. 1e and f). The color gradient in Fig. 1e indicates the strength of coupling in red and blue and conservation in green. The darker the color is, the stronger the coupling is (or the higher the conservation is), revealing that residues have stronger internal interaction in each community, but weaker couplings for residues from different communities. For instance, the MR1 provides a binding platform through the *α*_1_ and *α*_*a*_ domains in the MAIT *αβ*TCR^34^, characterized by communities in green and blue (Fig. 1e, left). The community comprising residues at the CDR loops of the V_*α*_ and V_*β*_ domains that involve in binding activities of TCRs^30,31^.

To understand how the G7 *γδ*TCR present a distinct binding strategy from others, we performed analysis of the residues communities of the MR1 domain. The community in red comprises a set of amino acids on the *α*_2_ domain that makes up the groove for holding the V_*γ*_ and V_*δ*_ domains (Fig. 1f, right). The residues not only provide physical contacts for the antigens but also act as a hub to mainly anchor the V*δ* domain. The residues constitute the community in blue, mainly located on the *β* -strand on the *α*_3_ domain. It provides direct interaction between the *α*_3_ domain and the V_*δ*_ domain, as the identified residue, E259, directly interacts with the residue, W105, at the interface of the V_*δ*_ domain. By superimposing the two MR1 proteins, the side-chain of the residue E259 present almost 180° rotation, by comparison to that in the MR1 of the MAIT *αβ*TCR-MR1, to form electrostatic attraction (Fig. 1f, top right), suggesting that it would be critical to contribute the unexpected docking topology that binds the underneath of the MR1 antigen presenting groove.

### 2.2 Functional-relevant residue communities

The *γδ*TCRs have a distinctive T-cell receptor that consists of one *γ* chain and one *δ* chain, but it still remains largely unknown how the antigenic molecules trigger *γδ*TCRs for signaling. As revealed by structural studies, CDR loops are functionally significant for antigen recognition in antibodies and the *αβ*TCRs^35–38^, as the docking mode between the TCRs and the ligands depends on the CDR loops, using CDR1 and CDR2 loops for contacts around a central region, particularly, CDR3 loops present major binding interactions^34,35^. One distinct interaction of the *γδ*TCR is underneath of the MR1 platform, and the interaction was mediated by the V_*δ*_ 1 region, mainly dependent on the V_*δ*_ 1 CDR3 loop^29^. Although the facts that the CDR3 loops exhibit high flexibility in binding activity have been achieved on antibodies and TCRs^36,37^, conservation analysis on the loops is not sufficient to characterize a functional requirement of CDR loops for the *γδ*TCRs to contact both the MHCs and the presented peptides. Coevolution between non-covalent residues are crucially important indicators of protein-protein interactions, as it holds information for evolution and mediates essential biological phenomena. The TCR *γ* and *δ* chains likely depend on a nuanced strategy of binding at different interfaces, and we suspected that the changes in the coevolution might contribute to the distinct binding mode. Evolutionary couplings derived from coevolution indicate patterns of functional correlations within the TCR *γ* and *δ* chains, and they may be general signatures adopt different functional roles in proteins. However, noisy signals that arises from lacking of diverse sequences are contained in the couplings. We performed the spectral analysis on the evolutionary couplings that can clearly separate functional correlations from background noises on the generated MSAs of the TCR *γ* and *δ* chains. Within either the TCR *γ* or *δ* chain, residues have strong intr-community couplings and weak inter-community interactions (Fig. 2a). Within the TCR *γ* chain, one of the three communities, community I, is quantitatively distinct and independent to the others, and residues in the community I form an interface with the V_*δ*_ chain, while part of residues in the larger community III (C_*γ*_) are to interact with those on V_*γ*_, importantly, the remain residue in the same community mainly locate at the CDR2*γ*, one of functionally significant CDR loops^29^ (Fig. 2a, top and b), although strengths of couplings in the community III are weaker than those in the community I.

**Figure 2.**
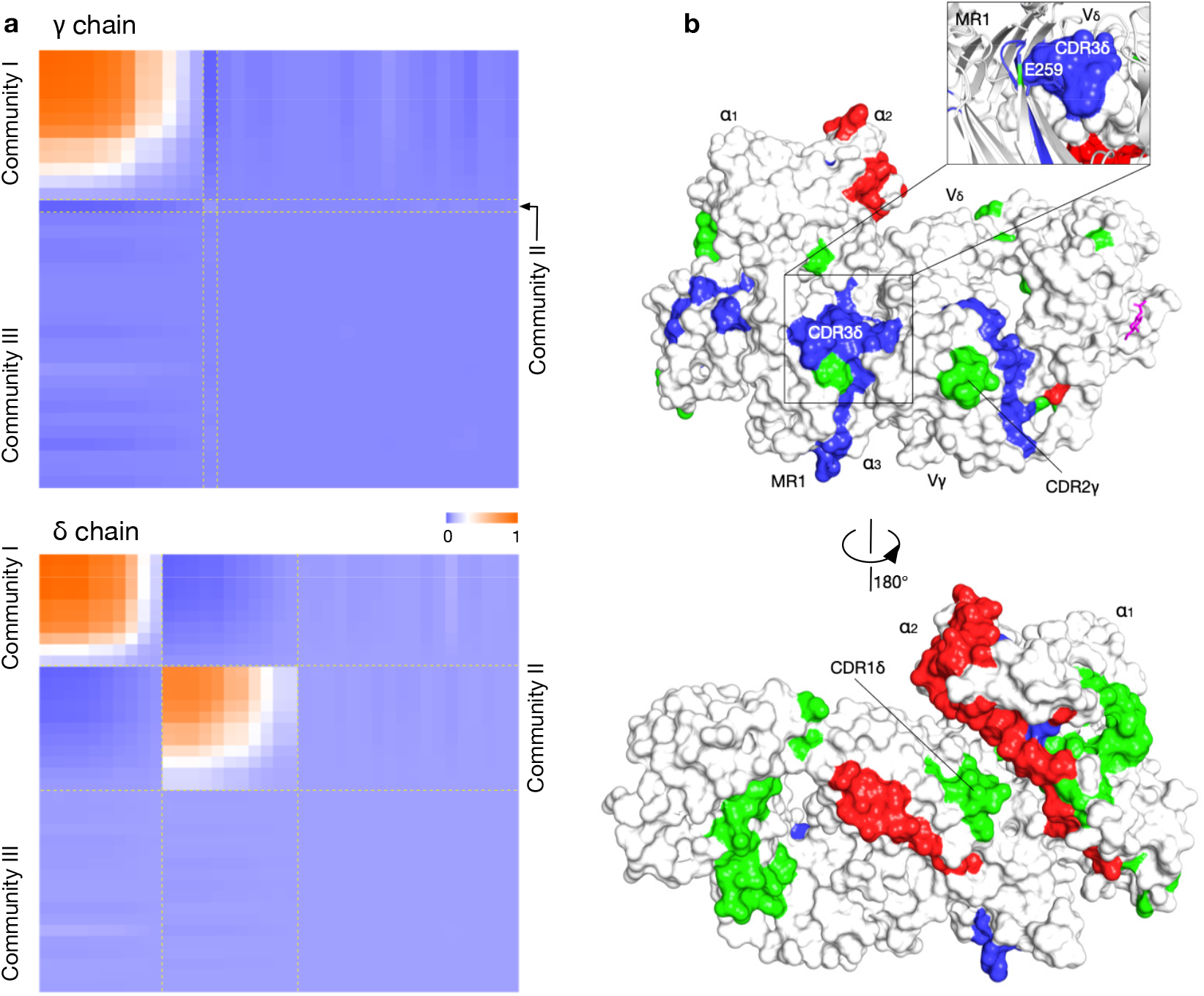
Molecular interactions. (a) Evolutionary signatures are revealed by highly ordered patterns formed by coupled residues. In each community, residues are sorted by descending based on their coupling strengths, displaying hierarchical patterns. (b) Residues in identifies communities located at functional sites. Left: the residues identified in different communities are colored red, green, and blue, respectively. Top: the identified glutamic acid (E259) seemly plugs into the interface of the TCR *δ* chain. Right: the CDR1*δ* was identified close to the interfaces between the MR1 and the TCR *δ* chain, and the identified regions on the *α*_2_ and *β*_2_M may provide the binding affinity to the TCR *γ* chain.

Fig. 2a, bottom, illustrates a distinctly hierarchical pattern between communities within the TCR *δ* chain. Residues in community I may have no important functional role for the binding mode in the TCR-MR1 complex, but those residues in the community II exhibited stronger correlations with the binding strategy, as the CDR3*δ* loop was identified at the interface of the *α*_3_ domain, suggesting the binding mode might be encoded during evolution. The three communities of residues were identified from the their couplings and mapped to the tertiary structures of the TCR *γ* and *δ* chains (Fig. 2b). The CDR1*δ*, CDR2*γ*, and CDR3*δ* were identified, whose functional importance consists with experimental findings^39^, particularly, the CDR3*δ* directly interacts with the *α*_3_ domain through the electrostatic attraction between the residue E259 of the *α*_3_ and W105 of the TCR *δ* chain. The residues in red of the *α*_2_ domain and those of the *β*_2_M chain form an interface that is close to the CDR1*δ*. Together, these theoretical analyses suggest that the distinct binding mode of the G7 *γδ*TCR might evolve from natural selection.

### 2.3 Flexibility of CDR loops

Proteins form protein-protein interfaces likely to stabilize binding modes. Accordingly, we investigated substitutions of amino acids to identify flexible regions that may decrease connectivity at interfaces. We computed differences between the substantive mutants and wild-type amino acids using the sequence statistical potential, and the difference (*E*_*mutant*_ *−E*_*wt*_) was measured as ‘energy’ to indicate stability and flexibility. The higher the energy is, the more flexible the position is.

As illustrated in Fig. 3, the TCR *δ* chain (top) exhibited more flexible positions by comparison with the *γ* chain (top) indicated by the energy differences. The CDR2*γ* was identified (community in green) at a flexible region of substitutions with lower energy than that of the wild-type, and another long loop that interacts with the *δ* chain also preferred more flexible. The substitution of residues in the identified community (blue, CDR3*δ*) presented with energies lower than zero, suggesting flexibility might allow rewiring connectivity between the *α*_3_ and the CDR3*δ*. For instance, position 259 of the *α*_3_ domain, which points into the pocket formed by the CDR3*δ* (Fig. 2b), is a glutaminic acid, but in the MAIT *αβ*TCR it has a rotated side-chain, which likely to sterically destabilize the binding affinity for the TCR *β* chain, perhaps partly accounting for the fact the docking topology of the G7 *γδ*TCR bind the underneath of the MR1 antigen presenting groove. To identify sequence changes sufficient to introduce binding affinity, we mutated the position in the TCR *δ* chain to alanine based on the findings (Fig. 2b). We evaluated the mutant by predicting its 3D structure using AlphaFold2-multimer^40^. We found that its affinity-alerting confirmed the ability to bind the MR1 antigen presenting groove. This highlights how the mutant that stabilizes an alternate strategy through the formation of an interface between the CDR3*δ* and the *α*_3_ domain can crucially influence their binding. Briefly, these findings present that those residue communities at interaction interfaces act together to drive the coevolution of the unexpected binding mode in the G7 *γδ*TCR.

**Figure 3.**
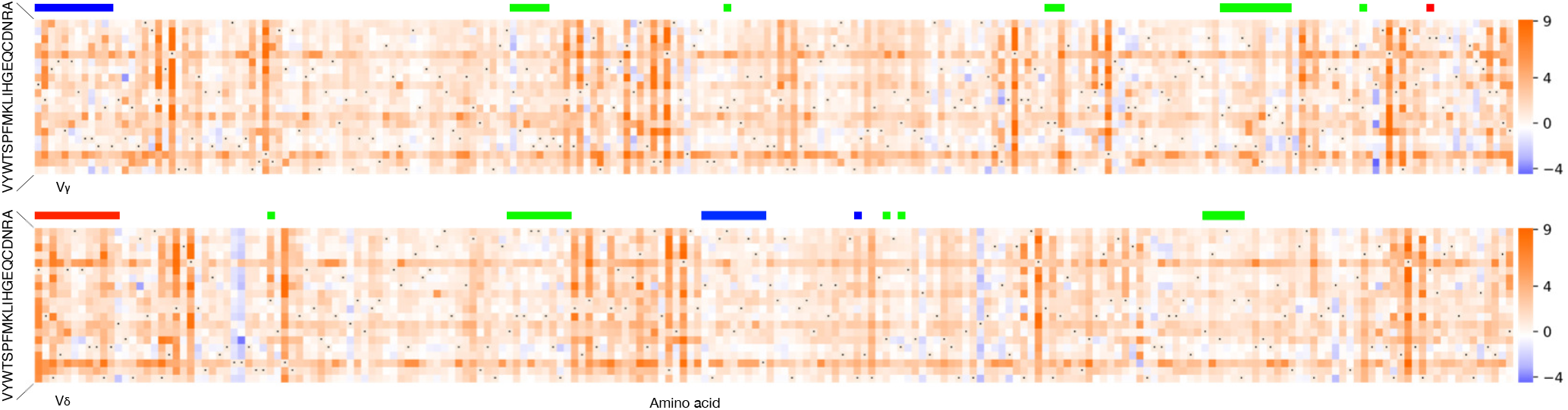
Substitutions of amino acids using the sequence statistical potential. The substitution matrix representations of the *γ* (top) and *δ* (bottom) chains, and the color bars on each representation indicate the regions of residues in the corresponding communities. The darker the orange is, the more favorable the substitution is preferred.

## 3 Discussion

In the present study, we develop a physics-informed approach–signatures of evolutionary couplings–to infer protein function at the residue level and leverage the model to produce function-related communities of residues for the G7 *γδ* T-cell reporter. The comprehensive residue communities in the present study provide a number of important insights at the residue level into protein evolution and function. First, functionally relevant communities are common and associated with different biochemical proprieties, suggesting residue communities as important signatures. Second, the identified residue communities present theoretical interpretations for the distinct mode of the G7 *γδ* T-cell reporter binding underneath the MR1 antigen presenting cleft, indicating that evolutionary information allows us to understand protein function from sequence alone. Third, residues in communities located in either protein cores or surfaces have a heavy effect on its stability and binding affinity, presenting a clue of high co-evolvability for molecular biases in TCRs and providing a basis for experimental investigations.

The primary advantage of our model is learning over evolutionary data to identify functional sites. Our method enables us to facilitate the rapid and comprehensive mapping of functionally important residues in highly ordered communities and evaluate mutants using the sequence statistical potential for diverse molecules, interactions and pathways, and our approach could be useful in evaluating residues for guiding experimental efforts in the future. Yet, there remains important challenges to capture disease-related variants. First, diseases may result from co-variants, but it remains speculative for different disease severity from the combinations of variants. Second, the present report does not cover the experimental effects in diagnostics or therapeutics, as our analysis is computational demonstration of how the evolutionary signatures shape proteins for biological activities.

## 4 Methods

### 4.1 From protein sequences to evolutionary information

Sequences of the MAIT *αβ*TCR-MR1-5-OP-RU and the G7 *γ* det TCR-MR1-5-OP-RU complexes were collected by searching each chain against the Uniclust30 databases^41^ (release Jun-2020) using HHblits (version 3.3.0)^41^ homology search tools with the default parameters at an E-value threshold 0.001. Following a filtering step, the generated sequence alignments were trimmed by two rules for satisfied coverage in sequences^33,42^: (1) a single site with more than 90% gaps across the MSA will be removed; and (2) a sequence with the percentage of gaps more than a given threshold (80%) will be deleted from the MSA. Finally, we collected 1,802, 1,311, 1,275, and 1,151 sequences for each chain in the G7 *γ* det TCR-MR1-5-OP-RU complex, and 1,800, 1,169, 1,314, 1,258 sequences were obtained for the chains in the MAIT *αβ*TCR-MR1-5-OP-RU complex. In the following procedure, each sequence *τ* in the trimmed alignment was re-weight for sequence diversity. The weight *ω*(*τ*) of a single sequence is defined by the sequence identity *I* using the normalized Hamming distance *D*_*H*_(*τ, τ*_*j*_) between the sequence *τ* and all other sequences (Eq. (1)).

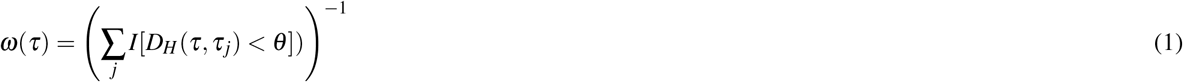

where *θ* is a threshold that controls the maximum diversity of pairwise sequences. In the present study, we used *θ* = 0.2 as default to weight the sequence for balancing diversity and conservation.

Given the MSA of *N* sequences by *L* positions for each protein, denoted as 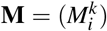, we can obtain an amino acid frequency at an individual position is 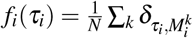, where *δ* = 1 if sequence *k* has amino acid *τ*_*i*_ at position *i*, otherwise *δ* = 0. Similarly, a joint frequency of amino acid between a pair of positions is formulated as 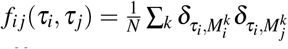.

The degree of conservation at each site was computed by Kullback-Leibler relative entropy^32^ as defined in Eq. (2),

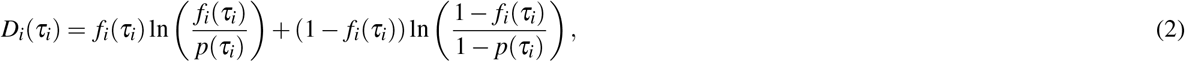

where *p*(*·*) is the background probability. The relative entropy *D*_*i*_ measures the degree of conservation of amino acids at a single position *i*.

Next, we inferred the couplings between pairwise residues from the same MSAs using the evolutionary couplings model^43^. The evolutionary process of a protein family can be mathematically modeled by a sequence generator at equilibrium. The generator produces a sequence *τ* with a probability *P*(*τ*) (Eq. (3)) from a distribution over the space of all possible sequences in the family.

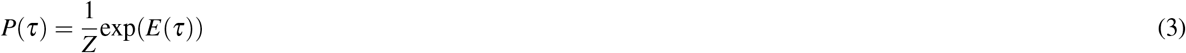

where *Z* is the partition function that normalizes the distribution by summing over the Boltzmann factors of all possible sequences in the family. The total energy (*E*(*τ*)) of the sequence *τ* is

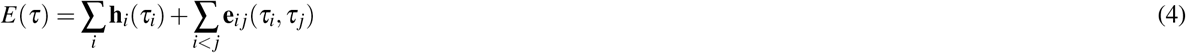

where **h**_*i*_ and **e**_*i j*_ are, respectively, site-specific bias terms of an amino acid and coupling terms between pairwise amino acids. The parameter **e**_*i j*_ measures the coevolving relationship between pairwise residues at two different positions of a MSA. Accordingly, **e** estimates evolutionary connections for all possible substitutions at two different sites, that is, a relationship matrix of *L×L×* 21 *×* 21, including 20 common amino acids and a gap, is to represent the connections of any pairwise residues at two sites in a protein sequence of *L*. Any correlation of pairwise residues of the sequence can be measured based on the matrix, termed sequence statistical potential. Given a sequence, its couplings matrix of *L × L* that describes evolutionary relationship of each pairwise residues is extracted from the relationship matrix.

### 4.2 Generating residue communities

Given the MSA of a protein family, the model parameters **h**_*i*_ and **e**_*i j*_ can be optimized by maximum likelihood. Here, we leverage a site-factored pseudo-likelihood approximation, instead of the full likelihood, to efficiently compute the partition function *Z* (Eq. 3), and the *l*_2_-regularization is to penalize the model to avoid over-fitting the sequences of the family. The objective function is defined

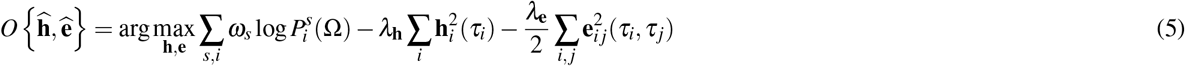

where Ω denotes all the sequences in the MSA, *ω*_*s*_ is weight of each sequence, the *l*_2_-regularization factors *λ*_**h**_ and *λ*_**e**_ are, respectively, set to 0.01 and 0.01 *· q ·* (*L −* 1), *q* is the total number of possible states. The conditional likehood 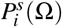 of sequence *s* at position *i* is defined

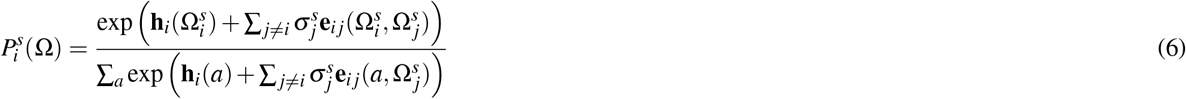

where 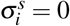 when the *i*th site is gapped, otherwise 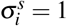.

To make the residue communities as much statistically independent as possible, the top three residue communities are defined based on two of the top five eigenvalues and their corresponding eigenvectors (**v**_*k*|*k*=1,*···*,5_) of the matrix **e**_*i j*_ (Eq. 4)^33^ as:

1. community I consists of residues at the *i*th position of 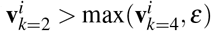;
2. community II includes residues at the *i*th position of 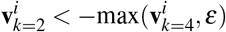; and
3. community III includes residues at the *i*th position of 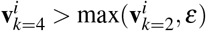.

We use an *ε* = 0.05 as a threshold to project the amino acids reduced from the coupling matrix and extract meaningful residue communities.

## Acknowledgements

The authors declare that this study received funding from Amoai Technologies Ltd, UK, which provided support for NJC and SYH. SYH is also supported by the Oxford Martin School and School of Systems Science.

